# Weather Characterization for Optimizing Genomic Prediction in Miscanthus Sacchariflorus

**DOI:** 10.64898/2026.03.18.712712

**Authors:** Ansari Shaik, Erik Sacks, Andrew D.B. Leakey, Hua Zhao, Jens Bonderup Kjeldsen, Uffe Jørgensen, Bimal Kumar Ghimire, Alexander E. Lipka, Joyce N. Njuguna, Chang Yeon Yu, Eun Soo Seong, Ji Hye Yoo, Hironori Nagano, Kossonou G. Anzoua, Toshihiko Yamada, Pavel Chebukin, Xiaoli Jin, Lindsay V. Clark, Karen Koefoed Petersen, Junhua Peng, Andrey Sabitov, Elena Dzyubenko, Nicolay Dzyubenko, Katarzyna Glowacka, Moyses Nascimento, Ana Carolina Campana Nascimento, Maria S. Dwiyanti, Larisa Bagment, Shatabdi Proma, Julian Garcia-Abadillo, Diego Jarquin

## Abstract

Environmental factors affect crop growth and development thus their consideration across sites and years become essential for genotypic evaluation. Genomic selection (GS) has been broadly implemented to accelerate breeding cycles by skipping field evaluations thus allowing early identification of outperforming genotypes. In this study, 7,740 phenotypic records corresponding to 516 *Miscanthus sacchariflorus* genotypes evaluated in five locations across three years were considered for analysis. Additionally, environmental data on six weather covariates was implemented to characterize similarities between locations. Different sets of locations of variable sizes were used for model calibration based on two cross-validations (CV00 and CV0) schemes leaving out one location at a time. Predictive ability across locations of the best model varied between 0.45 and 0.90 for both schemes. These results were compared to associate predictive ability in function of weather patterns between training and testing sets to allow model’s calibration optimization. We found it is feasible to optimize resource allocation by considering environmentally correlated sets. In most cases, the information from only one and, at most, two locations were enough to deliver better results than using all four locations, reducing training sets by up to 75%. The results obtained shed light on helping breeders make informed decisions considering weather data when designing evaluations.

## 1 INTRODUCTION

Plant breeding is a powerful tool implemented for developing traits for crop improvement to withstand biotic and abiotic challenges (Ceccarelli et al., 2010). Modern breeding techniques are key to delivering sustainable, profitable production of food, fiber, feed, fuel, and bioproducts (Moose & Mumm, 2008; Schmid & Thorwarth, 2014).

Emerging breeding techniques such as genomic selection (GS) (Meuwissen et al., 2001) and high-throughput phenotyping can help accelerate the development of new varieties, including adaptation to varying growing conditions across potential growing regions (Razzaq et al., 2021). GS is a marker-assisted tool that utilizes genomic-wide markers to predict the genomic estimated breeding value (GEBV) of the individuals and enhances the selection intensity in breeding programs (Goddard & Hayes, 2007; Meuwissen et al., 2001). Genetic gain can be significantly increased by making selections based on the GEBVs. In addition, the inclusion of genotype-by-environment interaction (G×E) in the analysis of multi-environmental trials is essential for identifying stable genotypes across sites (García-Arias et al., 2023).

A variety of approaches have been proposed to capture and explain genotype-by-environment interactions. Fisher and Mackenzie (1923) represented the differential responses of the genotypes to manure (management) by using a product instead of a sum, allowing capture non-linear interaction in potato varieties field trials. Sprague and Federer (1951) used variance components to partition the effects of genotypes, environments, and their interactions by equating the observed mean squares. Miller et al. (1962) found that the three-factor interaction (varieties × sites × years) is particularly important. Furthermore, factorial regression can be used on weather covariables to explain the genotype performance across environments (Vargas et al., 1999). Later, Jarquin et al. (2014) proposed the reaction norm model for the main effects of genomic, environment and weather gradients as a function of markers, location, and weather covariates, respectively, allowing the interaction between these factors. Several studies showed advantages of using models accounting for the interaction between the genotype and environment, but mostly in annual crops (Jarquin et al., 2014; Widener et al., 2021). *Miscanthus* acts as a potential perennial bioenergy fuel crop with a C4 photosynthetic pathway. It has shown environmental and ecosystem benefits by increasing soil quality and organic matter (McCalmont et al., 2017). Also, *Miscanthus* is gaining recognition due to its wider adaptability and higher biomass yield production with low inputs such as water, nitrogen, and heat. *Miscanthus* also achieves higher yield production and resource use efficiency than other bioenergy fuel crops (Chupakhin et al., 2021; Danalatos et al., 2007; Maleski et al., 2019).

Among various species of *Miscanthus*, *Miscanthus × giganteus* (M×g), a sterile allotriploid derived from the natural hybridization between *Miscanthus sinensis* (*M. sinensis*) and *Miscanthus sacchariflorus* (*M. sacchariflorus*), has shown high potential for biomass production (Anderson et al., 2011; Nishiwaki et al., 2011). Sheng et al. (2017) showed a remarkable heterosis of M×g for vegetative growth presenting more than 70 % yield increase than some accessions of the parental species. Breeding efforts of M×g are hindered by incompatibility between the parents, the sterility associated with the triploid genome, high establishment costs, and drought sensitivity (Clifton et al., 2019). In *Miscanthus* breeding, the most important trait to be improved is biomass yield (Njuguna et al., 2023). This trait is controlled by many factors such as physiological traits, weather patterns, resource availability (water, cold and heat), pests and diseases (Ings et al., 2013). Understanding the genetic diversity of the parental species of M×g is crucial for increasing biomass production and their adaptability to different environments and respective weather conditions (Njuguna et al., 2023).

As for other crops, the growing environment plays an important role in determining the biomass yield of *Miscanthus* (Arundale et al., 2014; Richter et al., 2008). In some sites, M×g suffers losses and damage during the first winter after establishment due to adverse weather conditions (Kalinina et al., 2017). In addition, cold temperatures in spring limit the leaf growth in *Miscanthus* (Farrell et al., 2006). Extreme weather events such as drought accelerate senescence in *Miscanthus*, inhibiting the mobilization of nutrients, while water logging stress promotes flowering in *M. sacchariflorus* leading to early flowering (Lee et al., 2018; Jensen et al., 2011).

Flowering timing is essential as it has an impact on crop productivity; early flowering reduces yields, and late flowering reduces winter survival (Dong et al., 2021; Jensen et al., 2011; Aurangzaib, 2012; Robson et al. 2012; Magenau et al., 2022).

The objective of this study is to test approaches for structuring training datasets for GS, which include information from different numbers of field-trial locations, as environmental data is leveraged in predictions of crop performance at unobserved location. This involved statistical analysis of phenotypic and genotypic data from a *M. sacchariflorous* population considering weather data collected from the NASA Earth Exchange (NEX) using the EnvRtype R package (Costa-Neto et al., 2021) for five locations observed during three years. The available weather information was used to determine environmental similarities between targeted locations (testing) and training sets to reveal the causes of the strong and weak association patterns between observed and unobserved sites. Due to the reduced number of locations, we conducted a weather characterization to find association patterns between locations and draw conclusions rather than include directly this information in the prediction models.

For this, three prediction models were considered, one based on phenotypic information (M1: S+T+L) and other two including genomic information as the main effect (M2: S+T+G) and in interaction with environmental factors (M3: S+T+G+G×S). These were evaluated considering two prediction problems of interest for breeders: CV00 (predicting untested genotypes in untested environments) and CV0 (predicting tested genotypes in untested environments) cross-validations scenario.

## 2 MATERIALS AND METHODS

### 2.1 Genomic Data

Previously, the genomic data was presented by Njuguna et al. (2023), considering a panel of 722 unique clonal genotypes of *Miscanthus sacchariflorus*. These were genotyped with single nucleotide polymorphism (SNPs) extracted restriction site-associated DNA sequencing (RAD-seq). Conventional quality control was applied where the SNP markers with around 50% of missing data and a minor allele frequency of less than 0.01 were eliminated (Głowacka et al., 2012). The resulting data consists of 34,605 SNP markers and 516 unique clonal genotypes available for analysis.

### 2.2 Phenotypic Data and Field Trials

For the 516 *M. sacchariflorus* genotypes, the phenotypic information consisted of 7,740 biomass yield records and for 15 other yield trait components collected in six trials established around the world in 2015 and scored between 2016 to 2018. The genotypes were observed in four temperate locations at higher altitudes – Sapporo, Japan, by Hokkaido University (HU); Urbana, IL, USA by the University of Illinois (UI); Chuncheon, South Korea, by Kangwon National University (KNU); Foulum, Denmark by Aarhus University (AU); and in one subtropical location at lower altitude – Zhuji, China by Zhejiang University (ZJU). Additional information about the *M. sacchariflorus* dataset can be found in Njuguna et al. (2023) and Widener et al. (2024).

### 2.3 Weather Data

The weather data was collected from the NASA Earth Exchange (NEX) using the EnvRtype package (Costa-Neto et al., 2021) in R software (R Core Team, 2024). This package allows access daily weather and elevation data using the “get_weather” function. It is a platform that assists in exploring and generating relevant meteorological information. The implemented dataset comprises information on six weather covariates: temperature mean [T2M], maximum [T2M_MAX], minimum [T2M_MIN], dew point temperature [T2MDEW], wind speed [WS2M], relative humidity [R2HM] collected at 2 meters, total precipitation [PRECTOT], all sky surface longwave [ALLSKY_SFC_LW_DWN], all sky surface shortwave downward irradiance [ALLSKY_SFC_SW_DWN]. In addition, these meteorological factors could be used to determine crop growth and development (Xu, 2016).

### 2.4 Statistical Models

Three prediction models (one phenotypic and two more including genomic data) were implemented in this study to evaluate the ability to deliver estimated breeding values (EBVs) under two cross-validation scenarios: (CV0), which mimics the problem of predicting tested genotypes in unobserved sites, and (CV00), which attempts to predict untested genotypes in unobserved environments. These models correspond to *i*) the main effects models based on site, harvest time and line effect (*M1: S+T+L*); *ii*) the genomic baseline STGBLUP (M2: *S*+*T*+*G*) model, which includes the genomic information via a covariance structure; and *iii*) an additional model including the genotype-by-environment interaction (M3: *S*+*T*+*G*+*G*×*S*). Detailed information about the models is mentioned below.

The BGLR package was used for fitting the models (Pérez and de los Campos, 2014).

#### 2.4.1 Baseline model

##### M1: S+T+L

Consider that 𝑦_𝑖𝑗𝑘_ represents the phenotypic response of the *i*^th^ (𝐿_𝑖_; *i* = 1,2,.., *n*) line observed in the site *j*^th^ (𝑆_𝑗_; 𝑗 = 1, …, 𝐽) measured at the *k*^th^ (*T_k_*; *k* = 1, 2, 3) time for dry biomass yield with 𝑆_𝑗_, 𝐿_𝑖_, and 𝑇_𝑘_ following identically and independent normal distributions (*IID*) centered on zero and respective variance components 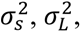 and 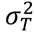 for site, line and time such that 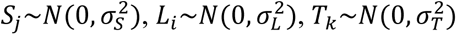 and µ being a common effect. *T_k_* effect of harvesting year allows the borrowing of the information within each site at different years. Additionally, the random residual term (𝜀_𝑖𝑗𝑘_) accounting for the unexplained variability follows a normal distribution centered on 0 and variance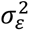 such that 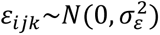. The resulting linear

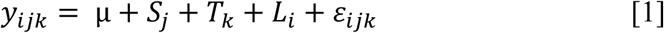

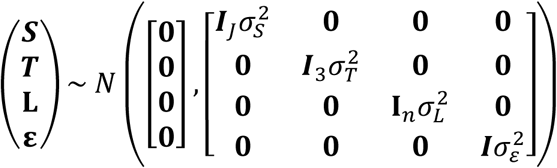

The limitation of this model is that the line (𝐿_𝑖_) effect, which only depends on phenotypic information does not allow the borrowing of information between the tested and untested genotypes. Thus, it is impossible to predict accurately unobserved genotypes. To address this inconvenience, genomic information could be used to connect training and testing sets.

#### 2.4.2 Environment and genotype main effects

##### M2: S+T+G

When the genomic information is available, the effect of the *i*^th^ genotype g_𝑖_ can be modeled as a linear combination based on the regression on *p* marker covariates and their corresponding marker effects such as 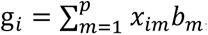, where 𝑥*_im_* represents the genotype of the *i*^th^ individual at the *m*^th^ marker position and 𝑏_𝑚_ is its corresponding effect. The marker effects are modeled as *IID* random variables (Habier et al., 2007; VanRaden, 2008) following normal densities centered on zero with a common variance 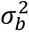, such that 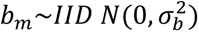. Hence, the vector of genomic values 𝐠 = {g_𝑖_} (*i*=1,2,…,*n*) follows a multivariate normal distribution with mean zero and covariance matrix given by **G** such that 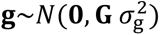 Here, **G** denotes the genomic relationship matrix calculated by 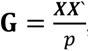, where 𝑿 represents the standardized (by columns) genomic matrix and the other model terms are as previously described. Hence, the linear predictor becomes

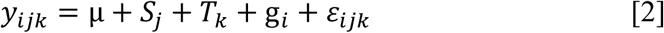

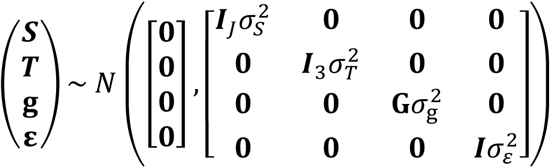

The GBLUP (Genomic Best Linear Unbiased Predictor model) considers constant genetic effects across locations impeding specific responses in locations/sites.

#### 2.4.3 Main effect and genotype-by-environment interaction

##### M3: S+T+G+G×S

The previous model accounts only for the main effects of markers (**g**) and sites (***S***), hence to capture the specific response of genotypes in sites, the interaction between the marker and the sites is included in model M3. This model term accounts for the reaction between the *i*^th^ genotype at the *j*^th^ site. Here, the interaction between the genomic effects and the sites are incorporated via variance-covariance structures such that 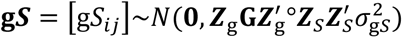 where 𝒁_g_ and 𝒁_𝑆_ are the incidence matrices that connect phenotypes with genotypes and sites respectively, and 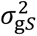 is the corresponding variance component. Additionally, the Hadamard product or the shur product (“o”) represents the cell-by-cell product or element-wise multiplication between two matrices of the same dimension (Styan, G.P, 1973). The resulting linear predictor is

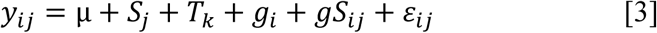

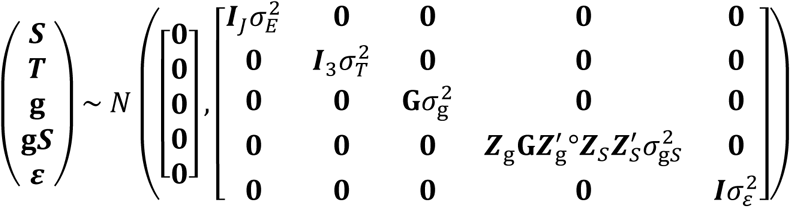

This model allows the borrowing of information between observations that are similar at the genomic and environmental levels.

### 2.5 Cross-validation

To evaluate the prediction accuracy of the models, two elaborated leave-one-out location cross-validation scenarios mimicking realistic prediction scenarios were considered (Jarquin et al., 2017; Persa et al., 2021). These cross-validation scenarios involve splitting the data into training and testing sets based on the planting locations. Then, comparisons are conducted to evaluate how well environmentally correlated locations based on a weather characterization help to increase the predictive ability of unobserved sites. To assess the usefulness of correlated locations to conduct predictions, different manners to compose training sets based on groups of locations of variable sizes (1-4) were considered similarly to Widener et al. (2021). These authors considered all combinations for different numbers of environments as training sets for predicting the trait performance of an annual crop, such as soybeans, for an unobserved testing environment.

In this study, a population of 516 unique clonal genotypes from *M. sacchariflorus* was used for the analysis where dry biomass yield was recorded across five different locations over three years. Here, one-third of the genotypes (172) was used as a testing set (the same across locations and years), and the remaining two-thirds (344) served for model training. Depending on the cross-validation scenario CV00 or CV0, the genotypes in the training set varied. For CV00, the remaining 344 genotypes observed in other environments and years were considered for model training in an attempt to mimic the prediction of novel genotypes in untested locations across years. To avoid unrealistic situations under the CV0 scenario, such as using data from the future (e.g., year 3) to predict the past (e.g., year 1), within the same location, the phenotypic records of the same genotypes were masked as missing values across years. Shaik et al. (2026) proposed a series of cross-validation schemes for implementing genomic prediction in perennial crops. These authors show how to design cross-validation schemes for avoiding contamination of the training set when considering locations with multiple years of information.

In addition, a modified version of the leave-one-out cross-validation (LOOCV) was implemented to conduct the model’s assessment under CV0 and CV00 scenarios. In general, the LOOCV iteratively uses all the units except the one (*s*-1) under validation for model training at different levels (e.g., phenotype, genotype, location, year, environment, etc.) For example, when predicting the performance of unobserved environments under the LOOCV, sequentially one environment is left out, and the remaining are used for model training. In this study, five locations (L_1_= AU, L_2_= HU, L_3_= UI, L_4_= KNU and L_5_= ZJU) were considered and the phenotypic information collected during the three years of evaluation at each of these are masked as missing values to conform to the testing set. Hence, in principle, the LOOCV involves iteratively training the models using information from four locations while the remaining is designed as testing or validation set. However, to evaluate the effects in predictive ability by considering correlated environments, different combinations of groups of locations considering between one to four were utilized, similarly as presented by Widener et al. (2021).

When predicting genotypic performance for one location, it is possible to choose from the remaining four locations subgroups of locations. For example, for a subgroup of only one location, a total of four unique combinations exist. Choosing two locations from the remaining four, it is possible to conform up to six unique combinations. Similarly, selecting three locations out of four produces a total of three combinations. Finally, there is only one single combination to select four locations. The total number of unique combinations is 15 (4+6+4+1). For example, predicting Location L_1_ we have the following combinations: one location ([L_2_], [L_3_], [L_4_], and [L_5_]); two locations ([L_2_, L_3_], [L_2_, L_4_], [L_2_, L_5_], [L_3_, L_4_], [L_3_, L_5_], and [L_4_, L_5_]); three locations ([L_2_, L_3_, L_4_], [L_2_, L_3_, L_5_], [L_2_, L_4_, L_5_], and [L_3_, L_4_, L_5_]); and four locations ([L_2_, L_3_, L_4_, L_5_]). Predicting the other locations, similar combinations are considered. Table 1 presents the 15 training combinations that can be formed for predicting a single environment. Hence, a total of 75 combinations (15 training combinations × 5 locations) resulted for predicting the five environments.

**Table 1.** Exhaustive combinations of groups of locations of different sizes for model calibration for one single location. Number of training combinations (right column) for a given number of locations (left column) considered as training sets. The number of possible combinations of size *k* to choose out of the four locations is given by 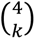.

| Location subsets | Number of combinations |
| --- | --- |
| 1 | 4 |
| 2 | 6 |
| 3 | 4 |
| 4 | 1 |
|  | 15 total possible combinations |

Considering all these cases helped to optimize the allocation of resources using correlated sites because they allow combinations to be found that effectively borrow information from one context to another more precisely to improve prediction accuracy. Combining the different cross-validation scenarios and the different manners to compose the training sets using groups of locations of variable size results in the following prediction strategies: CV00/groups and CV0/groups.

Figure 1 displays for the five different locations (panels 1A-1D) the 15 manners to compose the training sets based on groups of sites. For example, in panel 1A the objective is to conduct the prediction of the 172 genotypes that were not observed at any of the other locations. There, the first 4 columns represent the cases where only one location at a time was used for model training (L_2_, L_3_, L_4_, and L_5_). Then the remaining columns correspond to the six manners to choose two locations out of four, three out of four, and only one to combine all the remaining four (right extreme).

**Figure 1.**
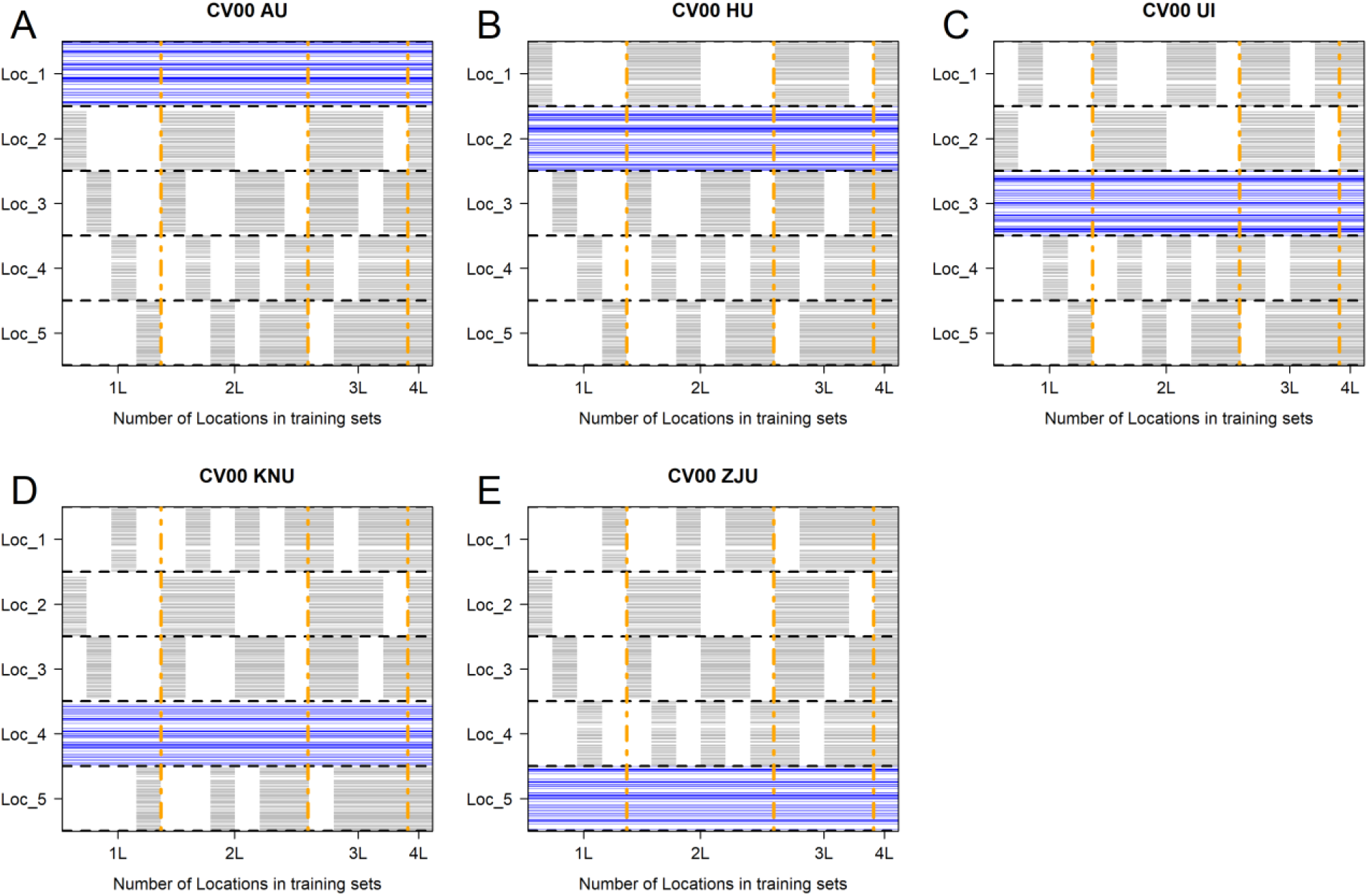
Graphical representation of prediction strategies based on CV00 cross-validation scheme (predicting untested genotypes in unobserved locations across years). The gray horizontal lines represent the two-thirds (344) of genotypes used for model training, the blue lines correspond to the testing set (174), and the columns within each panel indicate the different groups of locations (4 for 1L; 6 for 2; 4 for 3L; and 1 for 4L) used for model training for a total of 15. Panel A presents the prediction of Location 1 (L_1_ = AU) using the different location combinations (HU, UI, KNU and ZJU). Panel B shows the prediction of Location 2 (L_2_ = HU) using the remaining location (AU, UI, KNU and ZJU). Panel C shows the prediction of Location 3 (L_3_ = UI) by using the remaining location (AU, HU, KNU and ZJU). Panel D shows the prediction of Location 4 (L_4_ = KNU) by using the remaining four locations (AU, HU, UI and ZJU). Panel E shows the prediction of Location 5 (L_5_ = ZJU) by using the remaining four locations (AU, HU, UI and KNU).

Similarly to Figure 1, Figure 2 displays for the five different locations (panels 2A-E) the 15 manners to compose the training sets based on groups of sites. The only difference with respect to the previous case, is that some of the genotypes to be predicted in a testing location were probably observed already in other environments. Extra care was taken to ensure the phenotypes of the same genotypes were masked as missing values across the three years in the same location.

**Figure 2.**
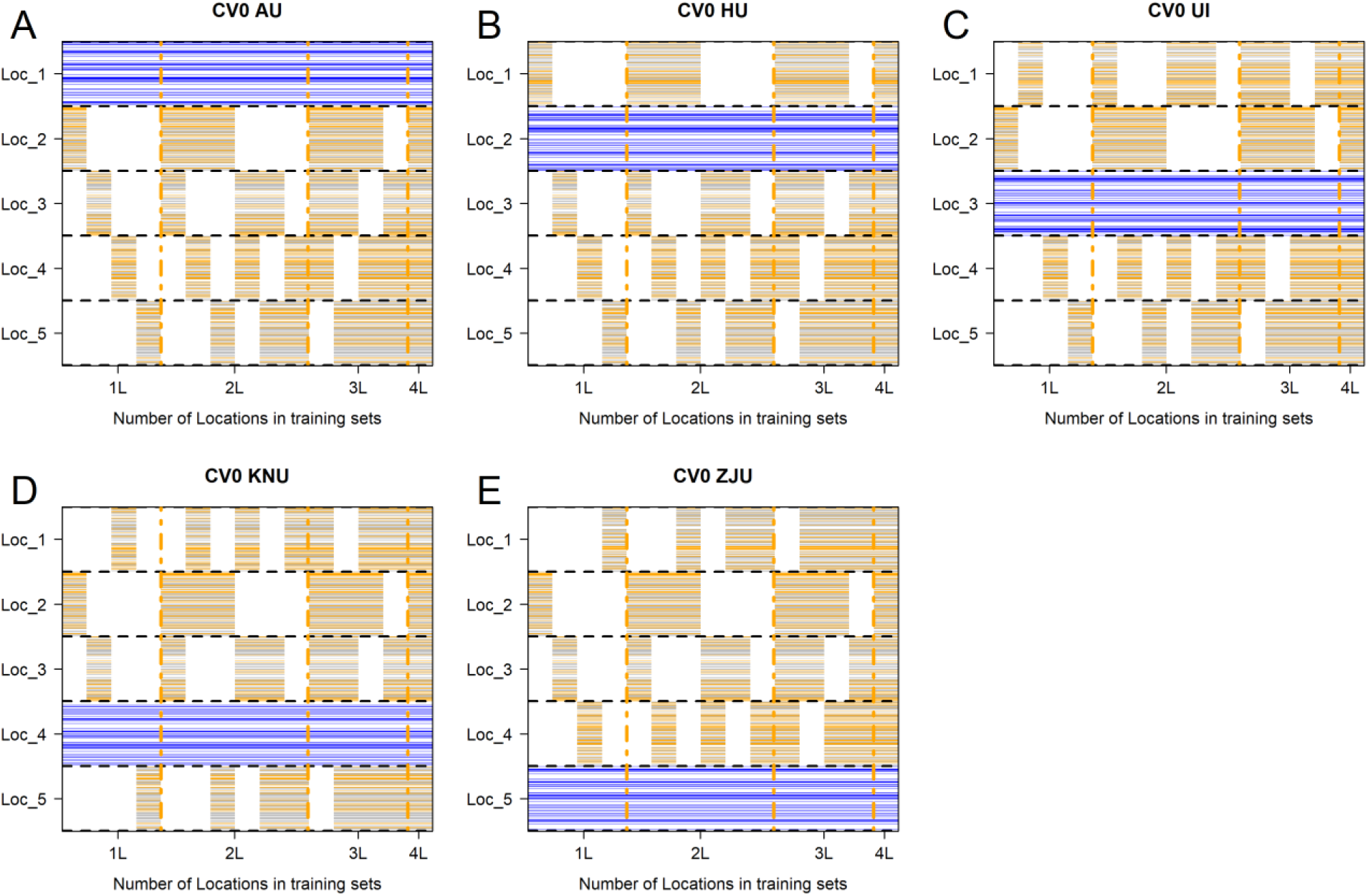
Graphical representation of cross-validation scheme (CV0. predicting tested genotypes in unobserved sites). The colored horizontal lines represent two-thirds (344) of genotypes for model training, the blue lines correspond to the testing set (174), and the columns within each panel indicate the groups of locations (4 for 1L; 6 for 2; 4 for 3L; and 1 for 4L) that were used for model training for a total of 15. Panel A presents the prediction of Location 1 (L_1_ = AU) using the different location combinations (HU, UI, KNU and ZJU). Panel B shows the prediction of Location 2 (L_2_ = HU) using the remaining Location (AU, UI, KNU and ZJU). Panel C shows the prediction of Location 3 (L_3_ = UI) by using the remaining location (AU, HU, KNU and ZJU). Panel D shows the prediction of Location 4 (L_4_ = KNU) by using the remaining four locations (AU, HU, UI and ZJU). Panel E shows the prediction of Location 5 (L_5_ = ZJU) by using the remaining four locations (AU, HU, UI and KNU).

## 3 RESULTS

### 3.1 Principal Component Analysis

The patterns of the nine environmental covariates across locations (L_1_-L_5_) and years (1, 2, 3) are presented in Supplementary Figures S1-S9. Clear trends for the different temperature covariates are observed (Figures S1-S4) between L_5_ and locations L_1_ and L_2_ with the first one being the warmest and the last two the coolest. Regarding the windspeed (Figure S6) L_1_ and L_5_ presented the lowest and highest speed. Relative humidity at 2 meters did not present clear trends in general; however, L_4_ presented the lowest values. Also, regarding total precipitation (Figure S7) there were no clear trends within and across years. For all sky surface longwave downward irradiance (Figure S8) L_2_ and L_5_ clearly showed the lowest and highest values. Finally, for all sky surface shortwave downward irradiance L_1_ presented the lowest values except in the middle of the period and L_3_ the highest values during the second half of the period. Since it is complicated to draw conclusions by considering the patterns of all variables at once, a PCA of the environmental kinship matrix 𝛀 was performed where 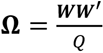, and ***W*** is the standardized (by columns) matrix of Q (9) weather covariates.

In general, principal component analysis (PCA) simplifies complex data by reducing its dimensions. PCA makes it easier to visualize patterns and clusters in data with many variables (Harrington et al., 2005; Abdi & Williams, 2010). Then, the eigenvalues and eigenvectors were computed using the *eigen* function of R-software. The outputs provide the eigenvalues and eigenvectors that quantify the variation of the matrix of environmental relationships (Abdi, 2007; Mishra et al., 2017) explained by the different components.

As a result, 15 principal components were produced representing the association patterns of the five locations over three years based on the collected weather information. The first principal component explains 62.6% of the variation in the data (see Figure 3A). In this case, the *x*-axis clearly separates locations L_1_ and L_2_ from L_5_, and L_3_ and L_4_ cluster together in the middle. This separation is in accordance with the different covariates based on temperature. The AU and ZJU locations have the coldest and warmest weather conditions, respectively. Location HU shows similar weather patterns to AU but warmer. UI and KNU are correlated sites and HU acts as a bridge sharing similar weather conditions. ZJU is not correlated with any of the sites. In addition, there is an interesting pattern connecting years from right to left. The second principal component (*y*-axis) accounted for 24.2% of the variation. In this case, L_1_ separates from L_3_ and L_4_ with L_2_ and L_5_ in the middle. This separation could correspond to differences in all sky surface shortwave downward irradiance.

**Figure 3.**
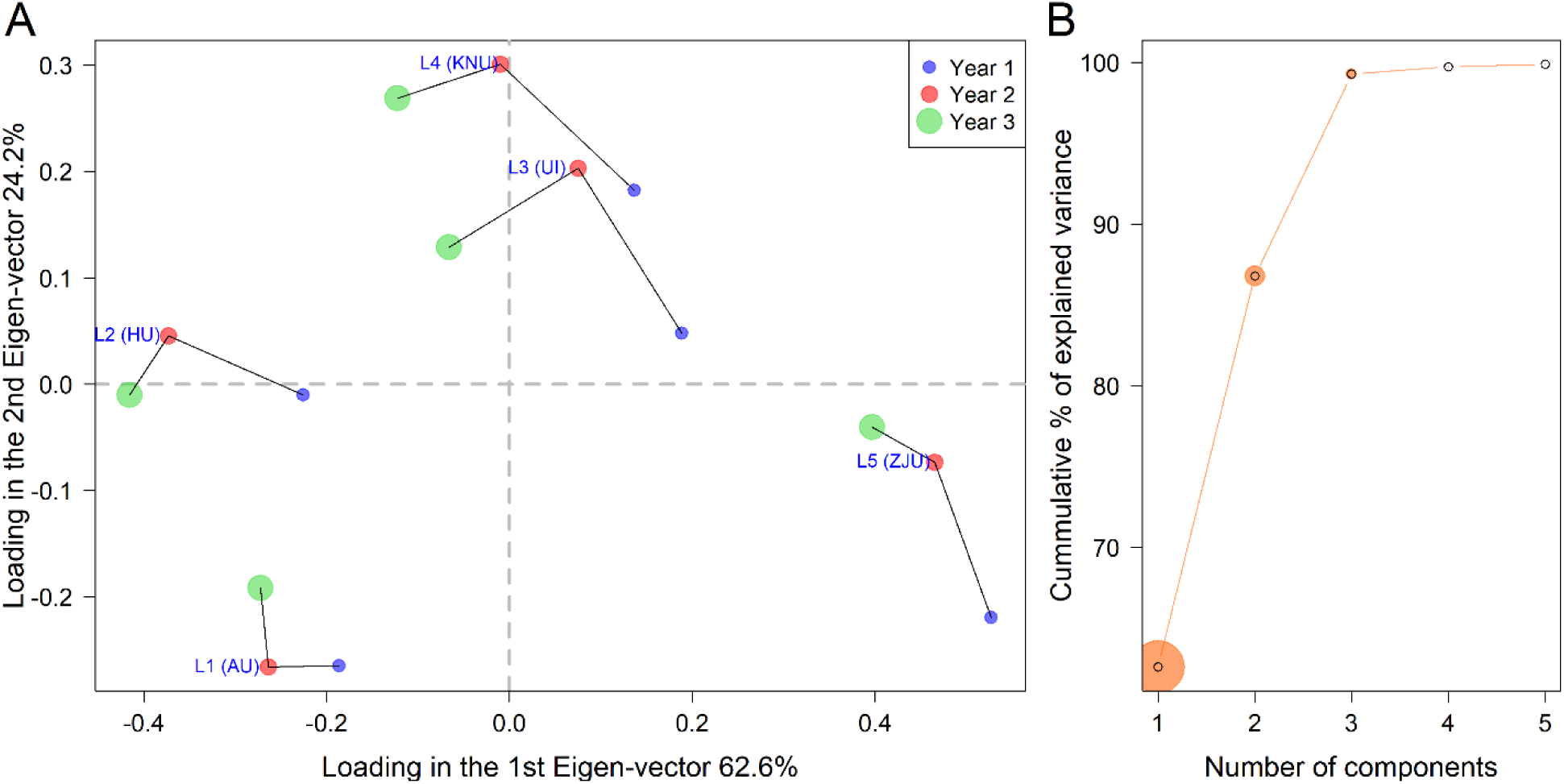
Environmental characterization using daily weather information on nine covariates (temperature mean, maximum, minimum, dew point, wind speed, and relative humidity at 2 meters, total precipitation, all sky surface longwave downward irradiance, and all sky surface shortwave downward irradiance) for three years (2016-2018) and five locations (L1 AU: Foulum, Denmark; L2 HU: Sapporo, Japan; L3 UI: Urbana Illinois, USA; L4 KNU: Chuncheon, South Korea; and L5 ZJU: Zhuji, China). (*A*) Biplot (*left panel*) loadings of two first eigenvectors. (*B*) Scree plot (*right panel*) cumulative proportion of variance explained by the principal components.

Figure 3B presents the scree plot of the eigenvalues, showing the cumulative proportion of variance explained by components. The *x*-axis presents the number of the principal components (1-5) while the *y*-axis shows the cumulative proportion of variance explained at each component. Combined the first and the second components explain close to 86% of the variance of the matrix of environmental relationships. Hence, these first two components provide enough information to conduct a weather characterization.

In Figure S10, for each location-by-year combination, the upper triangular matrix presents the phenotypic correlation between pairs of the location-by-year combinations. Similarly, the lower triangular matrix contains the sum of squares.

### 3.2 Assessment of Prediction Accuracy (CV00)

The performance of the models is evaluated based on prediction accuracy, measured as Pearson’s correlation between the observed and predicted values within the testing environment. This metric provides an indication of the model’s precision under the two considered cross-validation scenarios (CV00: predicting untested genotypes in unobserved locations across years, and CV0: predicting tested genotypes in unobserved locations across years) using the different training sets derived from the remaining locations.

CV00 scheme: Figure 4 presents the results for the five locations (panels A-E) and the three years for different sets of training locations of variable sizes and two different models. In this scenario, the baseline M1 (*S+T+L*) was excluded since its results are centered on zero, complicating the visualization of results of the models M2 and M3 due to small differences between these.

**Figure 4.**
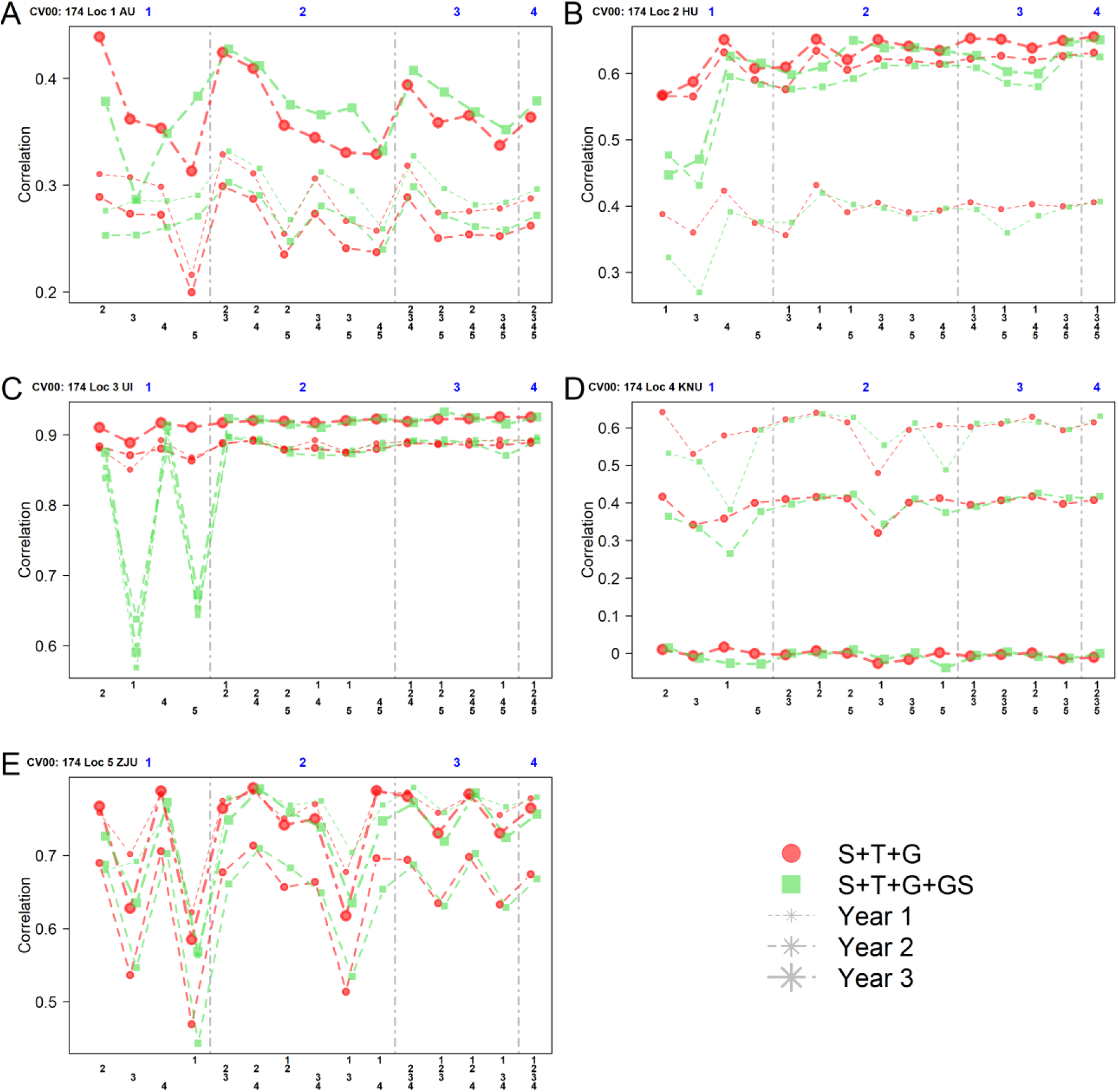
Predictive ability of dry biomass yield considering models M2 (*S+T+G*; blue color) and M3 (*S+T+G+G*×*S*; orange color) under the leave one location out cross-validation scheme for five locations and three evaluation years (L_1_ AU, L_2_ HU, L_3_ UI, L_4_ KNU, and L_5_ ZJU). Panels A, B, C, D, and E present the results of locations AU, HU, UI, KNU, and ZJU, respectively.

#### 3.2.1 Predicting L_1_ (AU)

For predicting dry biomass yield for the 172 genotypes in location 1 (AU), data from the remaining 374 genotypes observed in locations HU, UI, KNU and ZJU were used. Here, year 3 (L_1_) showed the highest prediction accuracy (Figure 4A), then year 1 and year 2. Predicting L_1_ in year 1, the two-location combinations {L_2_, L_3_} returned the highest correlation. In general, M2 and M3 performed very similarly, with the latter returning slightly better results. Similar results were obtained in year 2 where the most convenient strategy was also to consider only sets of only two locations for model training with the same locations as in the previous year {L_2_, L_3_} returning the best results. A slight reduction in predictive ability was observed compared to year 1, with models M2 and M3 returning similar results (∼0.30). In year 3, a relative improvement in prediction accuracy was observed (∼0.44) when considering the most successful prediction strategies (model and training set): M2 model and {L_2_} training set. Then, the second-best result (∼0.43) was obtained when considering two locations for training {L_2_, L_3_} for both models (M2 and M3). For this location, increasing the number of locations for model training did not improve predictive ability compared to more parsimonious sets.

#### 3.2.2 Predicting L_2_ (HU)

For predicting dry biomass yield in L_2_ (HU) across the three years (Figure 4B), the information from locations AU, UI, KNU and ZJU was considered. The results of this location were slightly higher compared to those from location 1 and ranged from 0.40 to 0.70. The prediction accuracy in year 2 and year 3 was better than in year 1. Predicting year 1 models M2 and M3 showed similar results, with {L_4_} and the combination {L_1_, L_4_} being the most convenient. Here, M2 achieved the highest prediction accuracies (∼0.43). In year 2, similar to year 1, {L_4_} and {L_1_,L_4_} again proved as the most effective combination when implementing model M2 (∼0.63). Whereas in year 3 the overall prediction accuracy improved compared to years 1 and 2. The four locations {L_1_, L_3_, L_4_, L_5_} performed slightly better when implementing M2 (∼ 0.66); however, similar results were observed when only location {L_4_} or the two locations {L_1_, L_4_} sets were used for model calibration. Across years, location {L_4_} consistently appeared when the highest correlation values were shown.

#### 3.2.3 Predicting L_3_ (UI)

Predicting dry biomass yield in L_3_(UI) consistently showed the highest correlations across years (Figure 4C) compared to the other locations. For predicting location 3 (UI), the information from locations AU, HU, KNU and ZJU were considered. Models M2 and M3 showed similar prediction accuracies in those cases where the information of two or more locations was used for model calibration. However, when using only one location as a training set, M3 showed the lowest predictive ability with locations L_1_, L_2_ and L_5_.

In year 1, different groups of locations of all sizes exhibited a robust prediction accuracy under M2, especially when location {L_4_} was involved. For example, M3 returned the highest prediction accuracy (∼0.90) with this single location. In year 2, the two locations {L_1_, L_2_} set returned the highest prediction accuracy (∼0.90) with models M2 and M3. Whereas in year the overall prediction improved compared to years 1 and 2. Here the three-location combination {L_1_, L_2_, L_5_} exhibited the highest prediction accuracy (∼0.93). However, these results were not different from those obtained when only one location was used for model training (e.g., {L_4_}). In general, considering {L_1_} consistently exhibited the lowest prediction accuracy across years.

#### 3.2.4 Predicting L_4_ (KNU)

Predicting dry biomass yield L_4_(KNU) the predictive ability declines year after year (Figure 4D). To predict L_4_ the information from the locations AU, HU, KNU and ZJU were considered. M2 and M3 exhibited similar prediction accuracies when two or more locations were used for calibration. In year 1 the highest predictive ability values were observed with M2 (∼0.64). The model performed similarly when one {L_2_}, two {L_1_, L_2_} or more locations were considered suggesting there are no improvements by including the information of additional locations. In year 2, a marginal decrease in prediction accuracy was observed compared to year 1. The three locations {L_1_, L_2_, L_5_} set returned the highest prediction accuracy (∼0.43) with model M3; however comparable results were obtained with both models using only one {L_2_} or two locations {L_1_, L_2_}. Whereas for year 3 a substantial decrease in prediction accuracy was observed for all groups of locations leading to null correlations.

#### 3.2.5 Predicting L_5_ (ZJU)

L_5_ (ZJU) is subtropical and has the most contrasting weather conditions compared to the other locations due to the highest temperature and relative humidity. Predicting dry biomass yield in L_5_ the information from the locations AU, HU, UI and KNU were considered. This location consistently showed a lower predictive ability (Figure 4E) when using data from {L_1_}. In year 1 models M2 and M3 showed similar accuracies for those cases where two or more locations were considered in training sets. Differences between models were observed when only one location was implemented, with locations {L_1_} and {L_4_} respectively returning the lowest and the highest values for this training set size. Overall, the two {L_2_, L_4_} and the three locations {L_2_, L_3_, L_4_} returned the highest prediction accuracy (∼0.79). Whereas in year 2, a slight reduction in prediction accuracy was observed compared to year 1, with M2 returning the highest value (∼0.71) with the two-location combinations {L_2_, L_4_}. In year 3 better results were obtained across training sets compared to year 2. Here again, the two locations set {L_2_, L_4_} returned the highest prediction accuracy (∼0.79). However, similar results were obtained with only {L_4_}. In general, across years, {L_2_} and {L_4_} consistently appeared when the highest correlation values were shown, in contrast to {L_1_} that returned the lowest values.

### 3.3 Assessment of Prediction Accuracy (CV0)

Figure 5 presents the results for the case when few genotypes in the testing set were also observed in those locations in the training set across the three years considering again groups of locations of variable size for model calibration.

**Figure 5.**
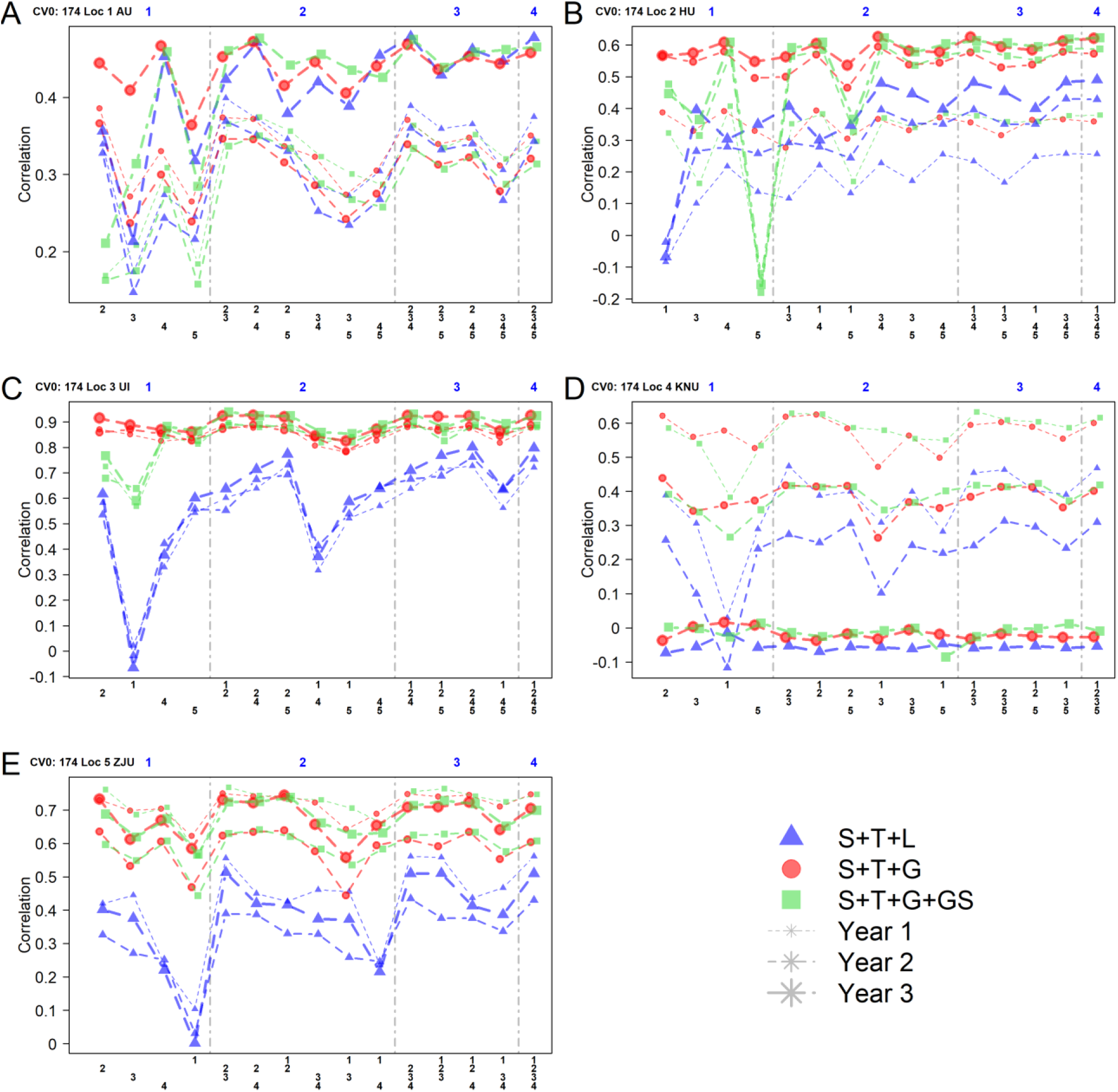
Predictive ability of dry biomass yield considering models M1 (*S+T+L*; grey color), M2 (*S+T+G*; blue color), and M3 (*S+T+G+G*×*S*; orange color) based on CV0 cross-validation (Predicting tested genotypes in unobserved sites) for five locations and evaluated for three years (L_1_ AU, L_2_ HU, L_3_ UI, L_4_ KNU, and L_5_ ZJU). Panels A, B, C, D, and E present the results of locations AU, HU, UI, KNU, and ZJU, respectively.

#### 3.3.1 Predicting L_1_ (AU)

Considering the results of the best model-training set strategies, year 3 exhibited a higher prediction accuracy compared to year 1 and year 2 (Figure 5A). In general, the baseline model M1, based only on phenotypic information, performed similarly to the models M2 and M3 when two or more locations were combined for model calibration. However, important differences were observed when only one single location was considered.

For year 1 the only one single location returned the best results with L_2_ (∼0.40). However, considering {L_3_} and {L_5_} results were negatively affected. This also was observed when considering groups of locations of size two or more. More specifically, the baseline model M1 returned the highest prediction accuracy (∼0.40) with the two-location combination {L_2_, L_3_}. Also, in year 2 these two locations again returned the best results, with model M1 delivering the highest prediction accuracy (∼0.37). Similar results were obtained with only {L_2_}. Whereas for year 3 this location showed the highest prediction accuracies when presented in calibration sets. Across years, slight improvements were observed when two or more locations were used in training sets for baseline model M1 and the interaction model M3. The baseline model M1 yielded the highest prediction accuracy (∼0.48) when the three locations {L_2_ L_3_, L_4_} were involved. However, considering only {L_2_} or {L_4_} also returned high results comparable to the best results from the other strategies involving more locations.

#### 3.3.2 Predicting L_2_ (HU)

The model’s prediction performance was the lowest in year 1, then improved in year 2 and year 3 (Figure 5B). In general, M2 and M3 outperformed M1. Across years, the single location {L_5_} consistently showed a negative correlation when it was considered as training set. In year 1 models M2 and M3 performed similarly considering two or more location combinations for model calibration. M3 returned the highest prediction accuracy (∼0.41) with the single location {L_4_} and these results were slightly better than those from M2 and when {L_1_} was utilized with either M2 or M3. Considering two or more locations the predictive ability was similar to the previous cases. While for year 2 the prediction accuracy consistently improved compared to year 1. The two locations group {L_3_, L_4_} returned the highest predictive ability (∼0.60). In addition, similar results were also obtained when three or more locations were used as training sets. Again, the single locations {L_1_}, {L_3_}, and {L_4_} returned comparable results to the higher correlations from other strategies for M2 only. In year 3 similar prediction patterns to the previous year were observed. Here, M2 returned the highest values, especially when single locations were considered. The highest predictive ability (∼0.63) was reached by models M2 and M3 with the two locations {L_3_, L_4_}. {L_5_} tends to reduce results when it was used alone or in pairs; however, with two or more locations the prediction accuracy slightly improved. Locations {L_3_} and {L_4_} consistently appeared when higher prediction values were observed.

#### 3.3.3 Predicting L_3_ (UI)

The results of predicting L_3_ (UI) are depicted in Figure 5C. Consistently these show higher and similar predictive ability patterns across years compared to other locations. In general, M2 and M3 showed similar results when a two or more-location strategy was used for model calibration. Differences were observed when only one location was considered. M1 returned the lowest values and showed null to negative correlations with {L_1_}. In year 1, models M2 and M3 returned similar results when two or more locations were considered. The model M3 exhibited slightly higher prediction accuracy (∼0.89) compared to other cases when the two locations strategy {L_1_, L_2_} was considered. Here, similar results were obtained with M2 and only any of these locations. In year 2, the three locations {L_1_, L_2_, and L_4_} set returned the highest prediction accuracy (∼0.90) with models M2 and M3. Similarly to the previous year, using M2, the information from only one location is enough to deliver results comparable to the more successful strategy. In year 3 a slight improvement in predictive ability was observed compared to years 1 and 2. Here, the interaction model M3 exhibited the highest prediction accuracy (∼0.94) with the two locations {L_1_, L_2_} strategy. Similar to previous years, good results were obtained considering only one location for model training with M2.

#### 3.3.4 Predicting L_4_ (KNU)

The results of predicting L_4_ are shown in Figure 5D. Interestingly, the predictive ability in this location declined over the years. The prediction accuracy in year 1 showed higher accuracies compared to year 2 and year 3. Year 3 showed null to negative correlations for all training sets and models. Overall, Model M2 and M3 outperformed model M1 in year 1 and year 2; however, in year 3, no differences were observed. Here, the highest correlations (∼0.64) were observed with M2 and M3 when two or more locations considered for model training. Again, the use of single locations, especially L_2_, returned similar values to the best results. For year 2 a reduction in prediction accuracy was observed compared to year 1. Here, M2 returned the highest prediction accuracy (∼0.44) when only L_2_ was considered. Regarding year 3, a significant reduction in predictive ability was observed. Besides year 3, {L_1_} alone or in pairs consistently showed the lowest results.

#### 3.3.5 Predicting L_5_ (ZJU)

As mentioned earlier, CV00 L_5_ (ZJU) is a subtropical environment and shows different weather conditions with warmer temperatures than the rest of location. Here, the baseline M1 model consistently showed the lowest prediction accuracy values across years and training sets (Figure 5E).

In year 1, M2 and M3 showed similar prediction accuracy with {L_2_, L_3_} returning the highest prediction accuracy (∼0.77). However, the differences compared to other training sets were minimal when considering models. Except when only {L_1_} was considered reducing the predictive ability. These two models performed similarly to the most accurate strategy when only location {L_2_} was used. In year 2, a slight decline in prediction accuracy was shown, with model M3 returning the highest prediction accuracy (∼0.64) with the two locations combination {L_2_, L_4_}. Here, again, the single location {L_2_} showed comparable results with model M2. In year 3, the two locations {L_1_, L_2_} set exhibited the highest prediction accuracy (∼0.74) with model M2. Comparable results were achieved by using the single location {L_2_}, and sets of two locations except for the set of locations {L_1_, L_3_} which reduced predictive ability. When considered individual locations for model training, {L_1_} continuously returned the lowest predictive ability across the years. However, when paired with other locations, the predictive ability slightly improved. Location {L_2_} frequently appeared when higher predictive values were observed.

## 4 DISCUSSION

The development of improved genomic prediction models requires the consideration of other layers of information such as environmental data to accomplish an optimum allocation of resources. Based on environmental factors, in this research we attempt to find association patterns between calibration and testing sets for predictive ability of dry biomass yield implementing genomic prediction models. This could lead to an optimization in the allocation of resources by requiring reduced groups of locations for model calibration for targeted locations. In most cases, we observed in many cases that one single location returns comparable results to considering all the four locations. Similar studies have been conducted but considering only annual crops such as soybeans (Widener et al., 2021). To our knowledge this is the first study implemented in perennial crops where larger impacts in genetic gains are expected due to long selection cycles.

Three prediction models were implemented on 7,740 dry biomass yield phenotypic records corresponding to 516 unique clonal genotypes of *Miscanthus sacchariflorus* observed during three years in five locations (516 × 3 × 5 = 7,740). Here, only one-third of the genotypes acted as testing sets in the predicted location. Depending on the cross validation scheme, calibration sets were composed from the remaining genotypes (CV00) or all of these (CV0). Determining informed manners to compose training sets for unobserved locations based on a weather characterization will help to optimize the allocation of resources while improving the predictive ability of dry biomass yield trait.

### 4.1 Assessment of Prediction Accuracy (CV00)

As mentioned before, the CV00 scheme represents the most challenging real-life prediction scenario. Here, the only way to connect the genotypes in training and testing sets is based on genomic information, while the environmental relationships between locations modulate the predictive ability.

#### 4.1.1 Predicting across years trait performance in L_1_ (AU)

Predicting the 172 genotypes in L_1_ (AU: Foulum, Denmark), the information from the remaining 374 genotypes evaluated in locations HU, UI, KNU and ZJU was considered. The correlations in L_1_ were found to be lower than in the other locations. Over the years, models M2 and M3 performed similarly. In year 3, model M2 showed the highest prediction accuracy (∼0.44), which represents an increase of 33% compared to year 1 (∼0.33) and 47% to year 2 (∼0.30). The model performed best when locations {L_2_} and {L_3_} were used as training sets (especially when these appeared together). In contrast, with {L_5_} the lowest values were obtained, except in year 3 where moderate results were shown. The results from the environmental time series (Supplementary Figures S1-S9) and the biplot (Figure 3A) suggest that location {L_5_} presents different weather patterns. Especially, showing a weak correlation with L_1_ (Supplementary Figure S10). On the other hand, training with location {L_2_} a highest prediction was obtained probably due to the moderate phenotypic correlation between these two locations (Supplementary Figure S10) and to the similar weather conditions, considering mean temperature, minimum temperature, dew point temperature and wind speed. When locations {L_2_} and {L_3_} were combined further improvements were obtained since these locations share more similar environmental conditions such as minimum temperature, total precipitation and dew point temperature with L_1_.

#### 4.1.2 Predicting across years trait performance in L_2_ (HU)

The prediction accuracy in L_2_ (HU: Sapporo, Japan) showed a slight improvement compared to L_1_ (AU) with models M2 and M3 presenting similar results. In year 3 the highest prediction accuracy (∼0.66) was obtained with the training set of four locations {L_1_, L_3_, L_4_, L_5_}, representing a relative improvement of 53% compared to year 1 (∼0.43) and 5% to year 2 (0.63) that implemented a two-locations {L_1_, L_4_} set for model calibration. This two-location combination constantly appeared when the higher predictive ability was observed. In addition, a moderate to strong phenotypic correlation (Supplementary Figure S10) was observed between these two locations with L_2_. More specifically, L_1_ showed similar weather conditions but it is slightly colder than L_2_. Based on the biplot and weather time series, L_2_ shares different weather conditions than L_3_ (UI) and L_4_ (KNU). Hence, borrowing the information from the remaining locations slightly improved the results.

#### 4.1.3 Predicting across years trait performance in L_3_ (UI)

Across locations, L_3_ (UI: Urbana Illinois, USA) showed the highest correlations. Over the years, M2 and M3 each presented similar correlations. In year 3, model M3 showed the highest prediction accuracy (∼0.93), which represents an increase of 3% compared to years 1 and 2 (∼0.90). Considering {L_1_} in calibrations sets consistently reduced predictive ability across years, mainly for M3. L_1_ presents lower temperatures than those from L_3_. Interestingly, the reverse case was different because predicting L_1_ with {L_3_} returned worse results. In both of these locations (L_1_, and L_3_) the interaction model M3 negatively affected predictive ability. {L_4_} showed a strong phenotypic correlation with L_3_ and also presented similar weather conditions (mean temperature, maximum temperature, minimum temperature and dew point temperature) in year 1. L_2_ (HU) slightly improves the prediction accuracy when combined with other locations by leveraging the information from the remaining sites.

#### 4.1.4 Predicting across years trait performance in L_4_ (KNU)

In L_4_ (KNU: Chuncheon, South Korea) there was a progressive decline in predictive ability over the years. In year 1, the highest prediction accuracy (0.64) was obtained by training the models with only one {L_2_} and two {L_1_, L_2_} location combinations while for year 2 (0.43) a decrease of 49% was observed. A more pronounced decrease was observed in year 3 (0.016). This significant reduction is probably due to an atypical year in the weather conditions for this location in year 3. In years 1 and 2, good to moderate results were observed for most of the training sets. Interestingly, when the information from only {L_4_} was used for predicting other locations, there were not observed drastically negative effects on predictive ability. Here, the best results were obtained when only location {L_2_} was involved, indicating a moderate to strong correlation with L_4_ (Supplementary Figure S10). These locations share similar weather patterns, including temperature covariates (Figures S1-S3). Whereas {L_1_} and {L_5_} presented more contrasting weather conditions (colder and warmer) compared to L_4_ (KNU) that falls in the middle. Lower predictive abilities were obtained when these locations conformed training sets.

#### 4.1.5 Predicting across years trait performance in L_5_ (ZJU)

Based on the weather covariates’ time series Location L_5_ (ZJU: Zhuji, China) presented the most contrasting environmental conditions (higher temperatures, relative humidity and dew point). In years 1 and 3, the highest prediction accuracy (∼0.79) was shown and it is 11% higher compared to year 2 (∼0.71). {L_1_} returned the lowest prediction accuracy across years (∼0.44 to 0.56) and a weak phenotypic correlation (Supplementary Figure S10) with L_5_. L_1_ and L_5_ are contrasting locations presenting respectively the colder and warmer weather conditions (mean temperature, maximum temperature and minimum temperature Supplementary Figure S1 panels A, B and C). This explains the low performance when these two locations alternated training and tested sets. {L_2_} and {L_4_} returned the highest prediction accuracy showing a strong environmental association with {L_5_} due to similar weather patterns for wind speed (Supplementary Figure S5).

### 4.2 Assessment of Prediction Accuracy (CV0)

The CV0 scheme represents the real-life scenario of predicting tested genotypes at unobserved sites. Here, some of the target genotypes to be predicted have been evaluated in other locations but not in the targeted location at any of the three years.

#### 4.2.1 Predicting across years trait performance in L_1_ (AU)

Predicting location L_1_ (AU: Foulum, Denmark) consistently showed lower prediction accuracy compared to the other locations, except for L_4_ in year 3. Within each year, all models performed similarly. Same as for CV00, the two-location {L_2_, L_3_} combination showed the highest prediction accuracy in year 3 (∼0.48), which is 30% higher compared to year 2 (∼0.37) and 20% to year 1 (∼0.40). The highest accuracy was achieved with the two {L_2_, L_3_} and three locations {L_2_, L_3_, L_4_} training sets. {L_2_} consistently improved the prediction accuracy due to similar weather patterns but slightly colder (Supplementary Figures S1-S3). Supplementary Figure S10 also shows a moderate phenotypic correlation between L_1_ and L_2_, L_3_, and L_4_; however, L_3_ and L_4_ varied from year to year. In summary, location {L_2_} alone was sufficient to provide accurate results and including L_3_ and L_4_ further enhanced predictive ability by borrowing the information between location with similar weather conditions considering maximum and minimum temperature, total precipitation, and dew point temperature.

#### 4.2.2 Predicting across years trait performance in L_2_ (HU)

Predicting L_2_ (HU: Sapporo, Japan) in year 3, M3 returned the highest prediction accuracy (∼0.63) representing an increase of 53% compared to year 1 (∼0.41) and 5% to year 2 (∼0.60). The best results were obtained when single location {L_1_} and {L_4_} and the two-location combination {L_3_, L_4_} were used for model training. Locations L_1_, L_3_ and L_4_ consistently appeared when the higher correlation values were observed probably due to similar weather conditions with L_2_. Weak to moderate phenotypic correlations were observed with location L_1_ (Supplementary Figure S10). However, this association varied from year to year, probably due to distinct weather conditions influencing distinct phenotypic responses.

#### 4.2.3 Predicting across years trait performance in L_3_ (UI)

L_3_ (UI: Urbana Illinois, USA) returned higher correlations compared to the other locations with M2 and M3 performing similarly when two or more locations were considered. In year 3 the highest correlation (∼0.94) was accomplished representing a slight increase of 4% compared to years 1 and 2 (∼0.90). The model performance was best with the two {L_1_, L_2_} and three-location combinations {L_1_, L_2_, L_4_}. A poor performance was observed only when the single location {L_1_} was used for model training.

#### 4.2.4 Predicting across years trait performance in L_4_ (KNU)

The prediction accuracy of location L_4_ (KNU: Chuncheon, South Korea) was found to decline over the years. Year 1 showed the highest prediction accuracy of (∼0.63) with the three-location {L_1_, L_2_, L_3_} set. The predictive ability was reduced by 30% in year 2 (∼0.44) with the single location {L_2_} returning the best result and finally by 97% in year 3 (∼0.016). The two locations L_2_ and L_3_ alone or in pairs showed the highest prediction accuracy, and these also shared similar weather conditions with L_4_. Furthermore, in this location the lowest accuracy was shown when using only using {L_1_} for model training. Low to negative correlations were observed in year 3, probably due to atypical weather conditions affecting production that year.

#### 4.2.5 Predicting across years trait performance in L_5_ (ZJU)

As mentioned earlier, location L_5_ (ZJU: Zhuji, China) belongs to a subtropical environment and shows contrasting weather conditions (warmer than other sites). L_5_ (ZJU) returned the highest prediction accuracy (0.77) in year 1 with M3, which was 20% higher compared to year 2 (0.64) and 4% than year 3 (0.74). Overall, results showed poor predictive performance when location {L_1_} was considered. The two location combinations {L_2_, L_4_} and {L_2_, L_3_} presented the best results. In general, L_2,_ L_4_, and L_5_ consistently showed moderate to strong phenotypic correlations.

Overall, CV00 and CV0 schemes showed similar results of predictive ability based on different approaches to composing training sets for predicting novel sites. Considering two or more locations for model calibration enhanced the predictive ability of the models predicting unobserved locations. Dealing with correlated locations, these returned the highest predictive ability due to homogenous weather patterns. Thus, genotypes performed similarly in correlated locations showing moderate to strong phenotypic correlations (Supplementary Figure S10). The lowest predictive ability of the models was observed when environmentally uncorrelated sites were considered for model training. Understanding the correlation among locations will help breeders by reducing phenotyping costs and minimizing field testing across various sites. Similar results were observed by Widener et al., (2021) in soybeans where these authors showed that a reduced number of locations is enough to ensure the highest correlations. In addition, these authors directly introduced the weather information in the models. However, due to the reduced number of environments it was not possible to appreciate significant improvements in predictive ability. In our case, dealing with a perennial crop we prefer to use the weather information to establish informed relationships between locations in training and testing sets. Shaik et el. (2026) introduced different cross-validation schemes including CV0 and CV00 considering a fivefold partition; however, these authors did not draw conclusions based on environmental relationships neither and/or to optimize the calibration sets based on groups of locations.

## CONCLUSIONS

Prediction models were implemented to assess different forecasting strategies for predicting variation in dry biomass production across a diverse population of *Miscanthus* genotypes in unobserved environments when considering the environmental variation present across the test sites. The leave-one-location-out-of-cross-validation (LOOCV) method was used to determine the most convenient way for composing training sets for each location using different groups of locations of variable sizes. In general, sets of reduced sizes (one or at most two) were sufficient to reach the highest correlations compared to the cases when all four locations were considered. Especially in those cases where environmentally correlated locations acted as training and testing sets.

Since the weather information was not explicitly incorporated in the models due to the low number of locations, considering the G*×*E interaction term did not improve the predictive ability compared to the main effects model M2 based solely on genomic information (M2). Our results identify that environmentally correlated locations help to improve prediction accuracy. The most accurate breeding values were achieved by the prediction models trained on only a single or a reduced group of correlated locations. The lowest predictive ability of models was observed with environmentally uncorrelated locations, whose did not share similar weather patterns.

Understanding how weather impacts *Miscanthus* crop across various environments is crucial for optimizing dry biomass production for biofuels and plays a main role in conducting the optimization of the allocation of resources. Exploring the weather conditions across locations would be helpful for breeders to reduce the phenotyping costs at multi-environmental trials and enhance the resource efficiency in the breeding process. Determining correlated locations with similar weather patterns will be helpful to better design experiments to conduct evaluations for selecting stable genotypes with high performance.

## Abbreviations

CV: Cross-validation
G×E: Genotype-by-Environment Interaction
GS: Genomic Selection
*Msa*: *Miscanthus sacchariflorus*
*Msi*: *Miscanthus sinensis*
*Mxg*: *Miscanthus giganteus*
PA: Predictive Ability

## Core ideas

The agriculture sector is adversely affected by severe weather conditions.

This study provides valuable insights to assist breeders in making informed decisions for trial evaluations.

Identifying the correlated locations with similar weather patterns can help designing better experiments and optimize the allocation of resources.

## ACKNOWLEDGMENTS

This work was funded by the DOE Center for Advanced Bioenergy and Bioproducts Innovation (U.S. Department of Energy, Office of Science, Office of Biological and Environmental Research under Award Number DE-SC0018420). Any opinions, findings, and conclusions or recommendations expressed in this publication are those of the authors and do not necessarily reflect the views of the U.S. Department of Energy.

## CONFLICT OF INTEREST

The authors declare no conflict of interest.

## DATA AVAILABILITY

The data sets and analyses implemented in this research can be found in https://doi.org/10.6084/m9.figshare.31787917

## SUPPLEMENTAL MATERIAL

**Figure S1.**
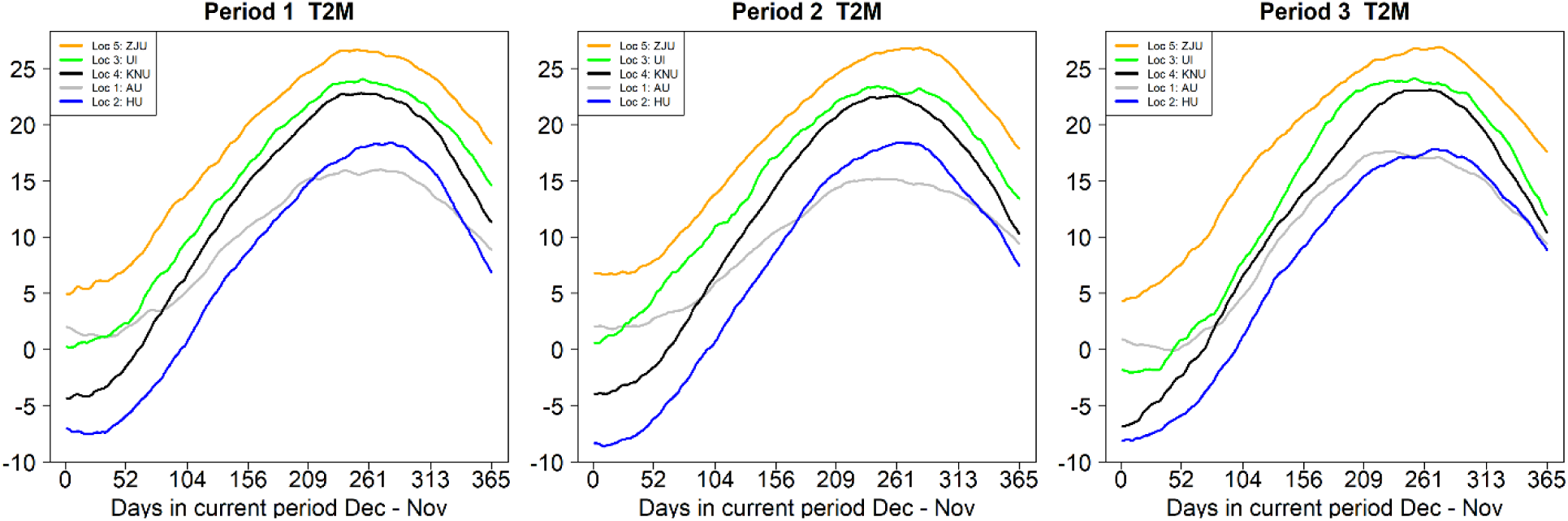
Time-series analysis derived for mean temperature at 2M (T2M) collected with the EnvRtype package for five locations during three years of *Miscanthus sacchariflorus* evaluations.

**Figure S2.**
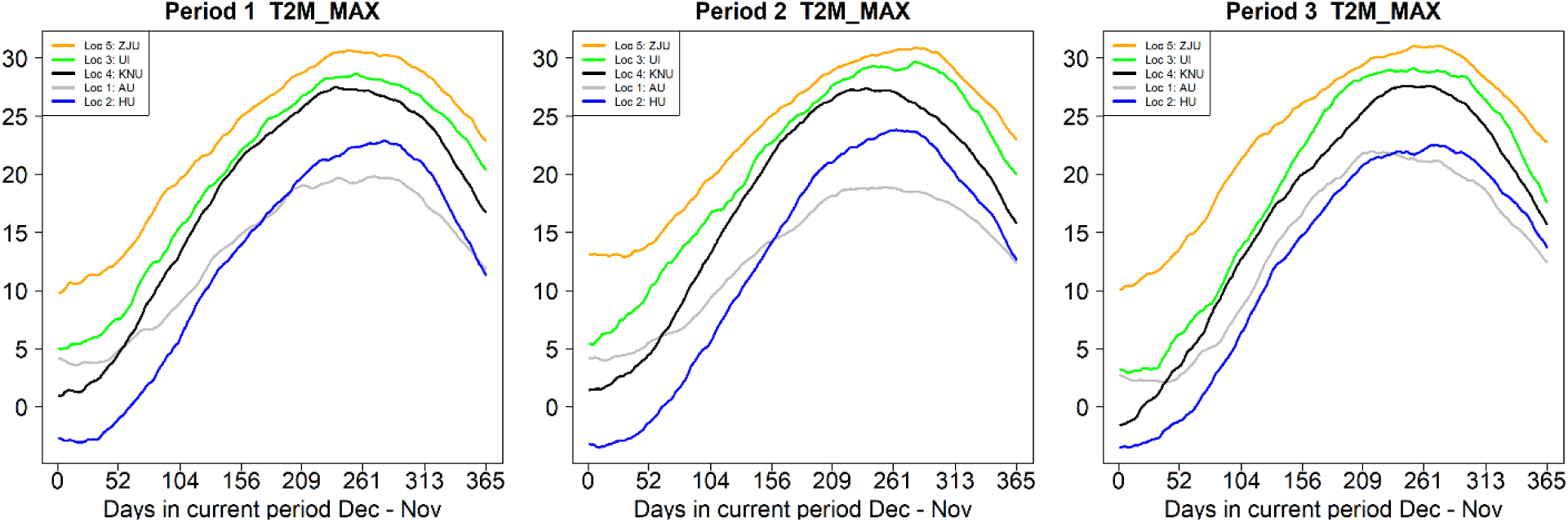
Time-series analysis derived for daily maximum temperature at 2M (T2M_MAX) collected with the EnvRtype package for five locations during three years of *Miscanthus sacchariflorus* evaluations.

**Figure S3.**
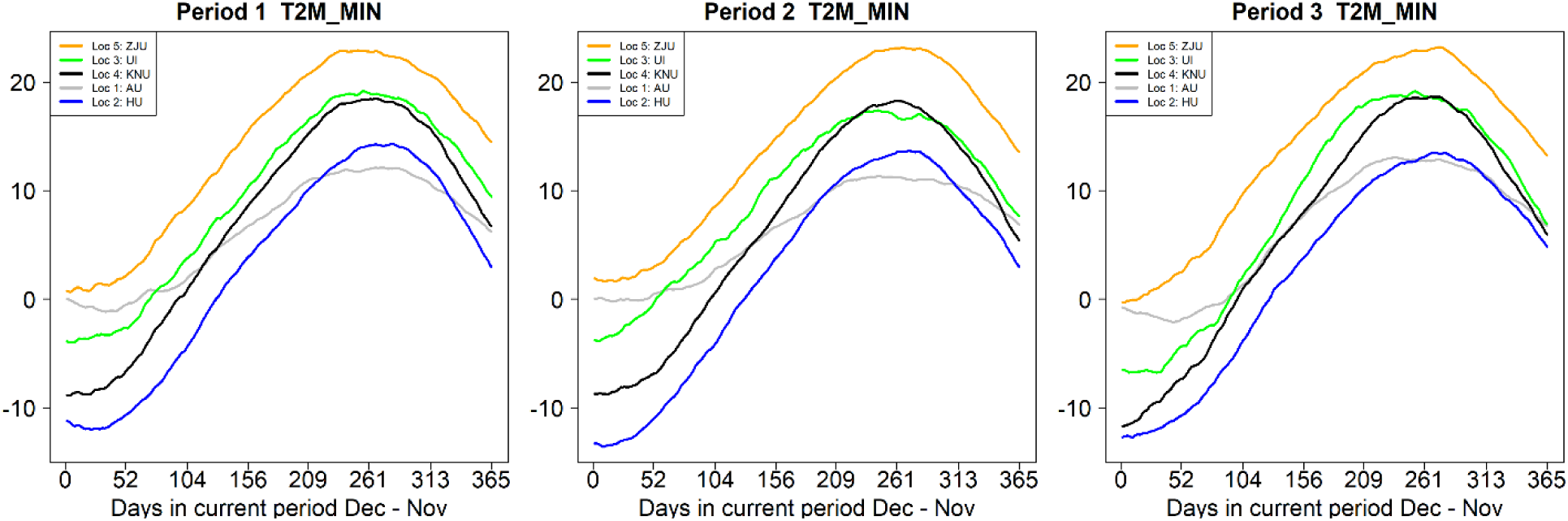
Time-series analysis derived for daily minimum temperature at 2M (T2M_MIM) collected with the EnvRtype package for five locations during three years of *Miscanthus sacchariflorus* evaluations.

**Figure S4.**
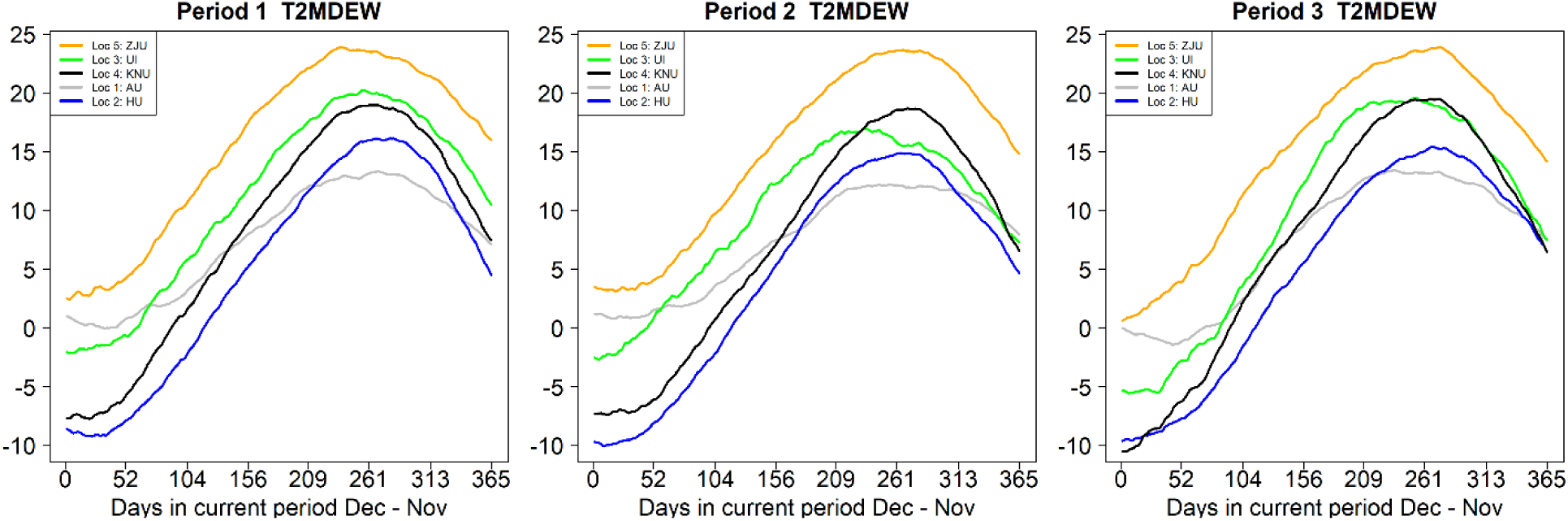
Time-series analysis derived for daily dew point temperature at 2M (T2MDEW) collected with the EnvRtype package for five locations during three years of *Miscanthus sacchariflorus* evaluations.

**Figure S5.**
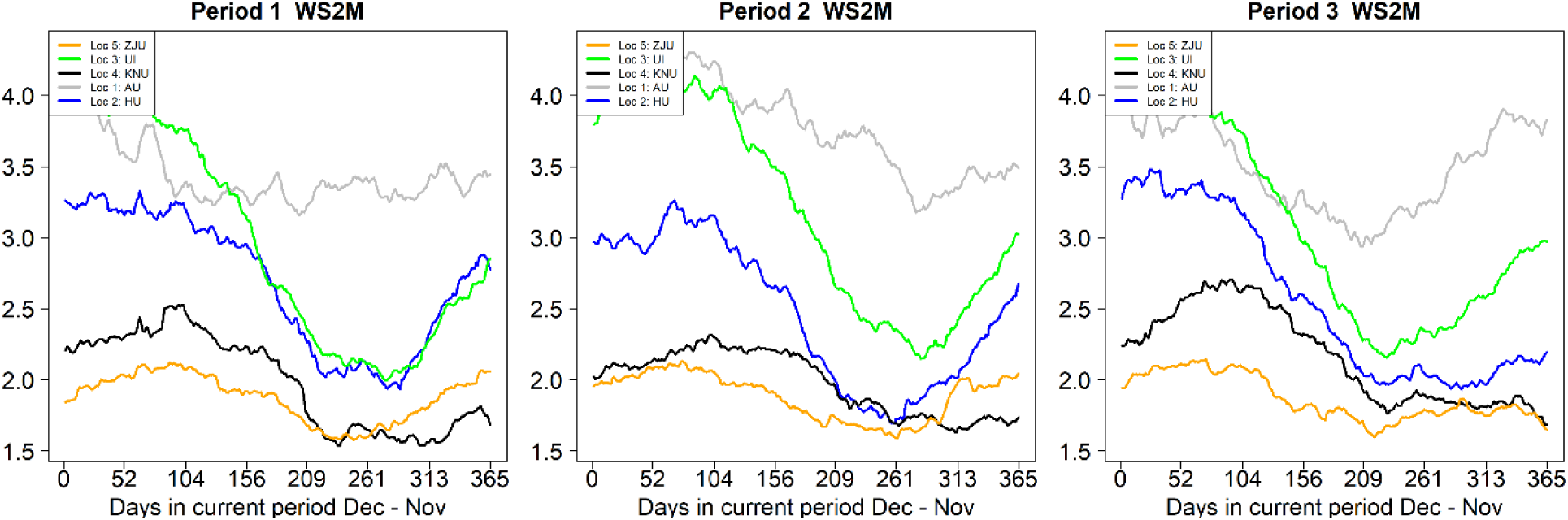
Time-series analysis derived for daily average wind speed at 2M (WS2M) collected with the EnvRtype package for five locations during three years of *Miscanthus sacchariflorus* evaluations.

**Figure S6.**
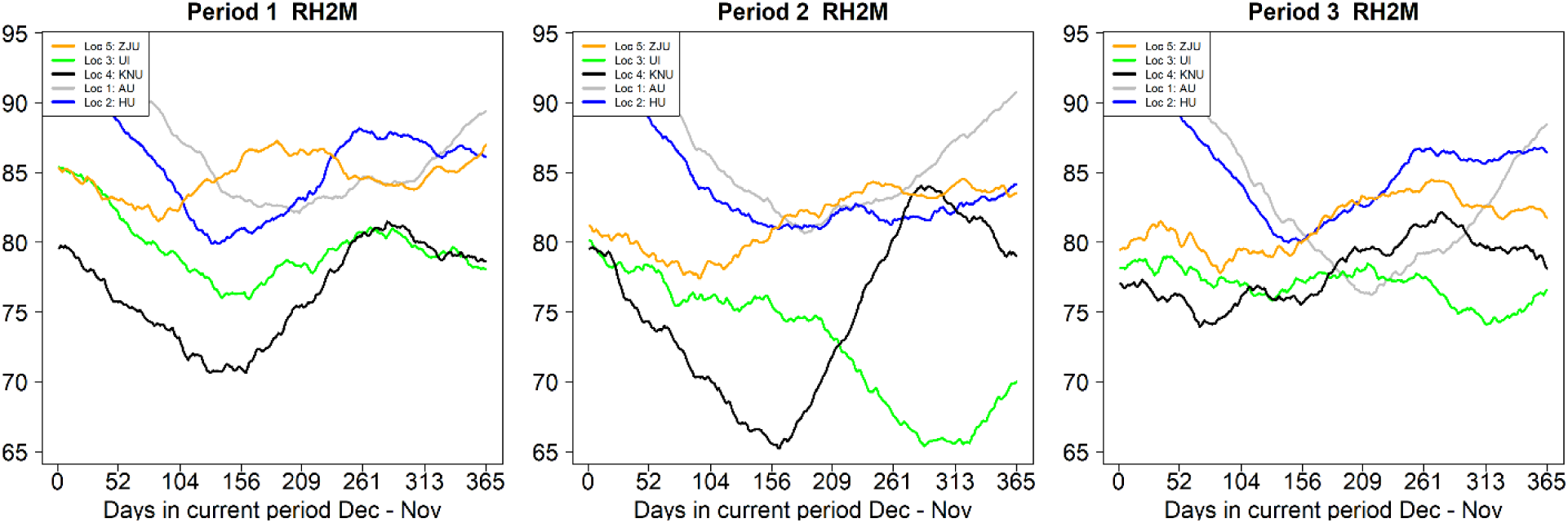
Time-series analysis derived for daily average relative humidity at 2M (RH2M) collected with the EnvRtype package for five locations during three years of *Miscanthus sacchariflorus* evaluations.

**Figure S7.**
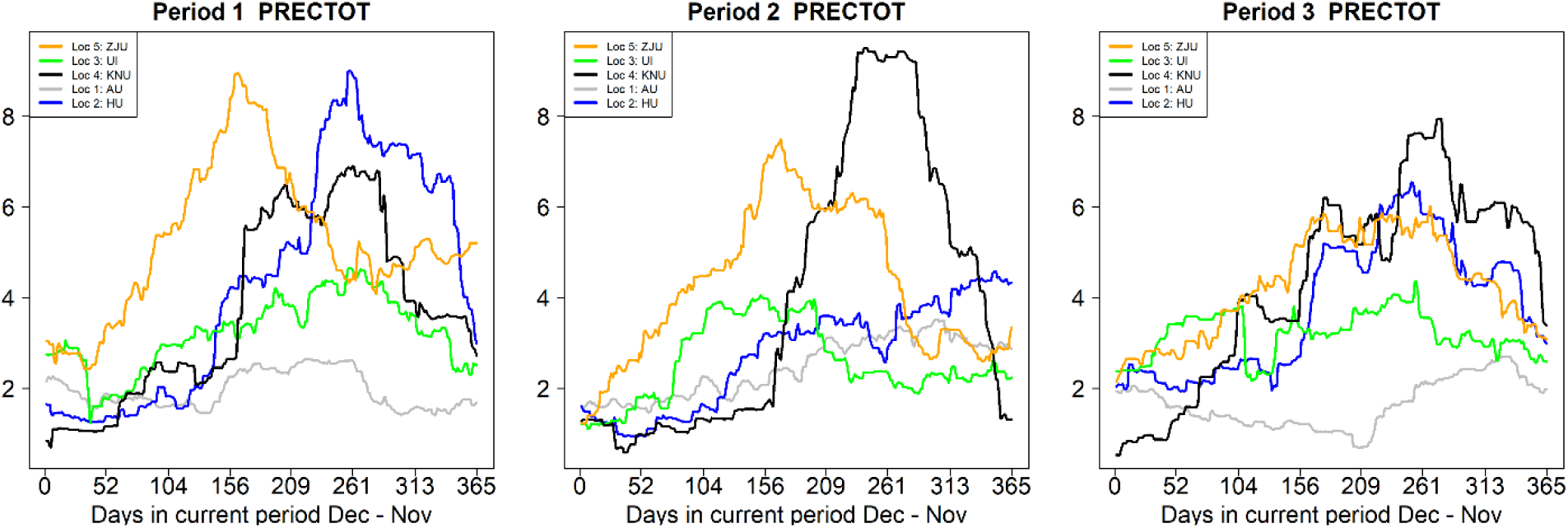
Time-series analysis derived for daily total precipitation (PRECTOT) collected with the EnvRtype package for five locations during three years of *Miscanthus sacchariflorus* evaluations.

**Figure S8.**
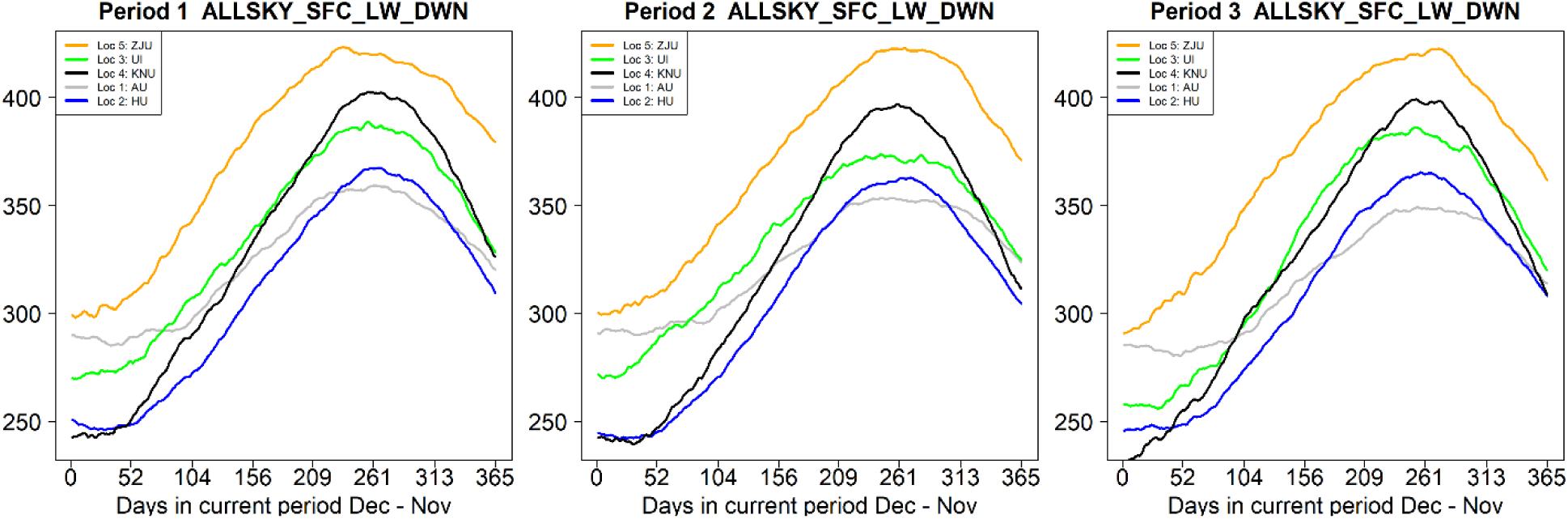
Time-series analysis derived for daily All Sky Surface Longwave Downward Irradiance (ALLSKY_SFC_LW_DWN) collected with the EnvRtype package for five locations during three years of *Miscanthus sacchariflorus* evaluations.

**Figure S9.**
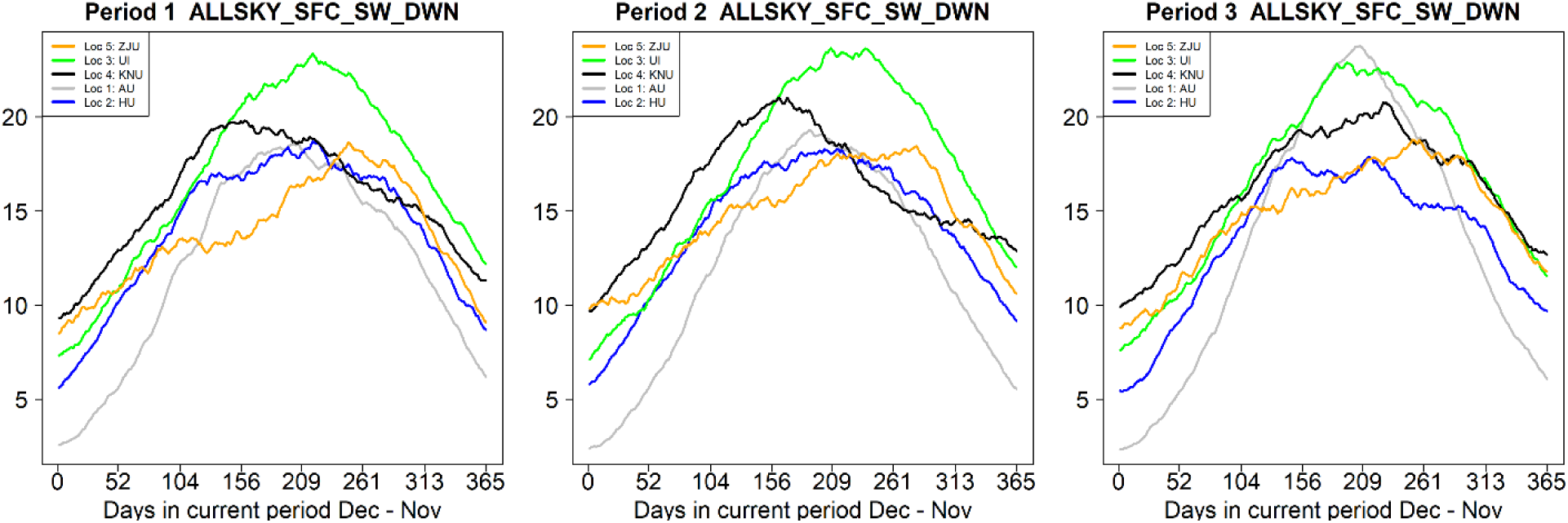
Time-series analysis derived for daily All Sky Surface Shortwave Downward Irradiance (ALLSKY_SFC_SW_DWN) collected with the EnvRtype package for five locations during three years of *Miscanthus sacchariflorus* evaluations.

**Figure S10.**
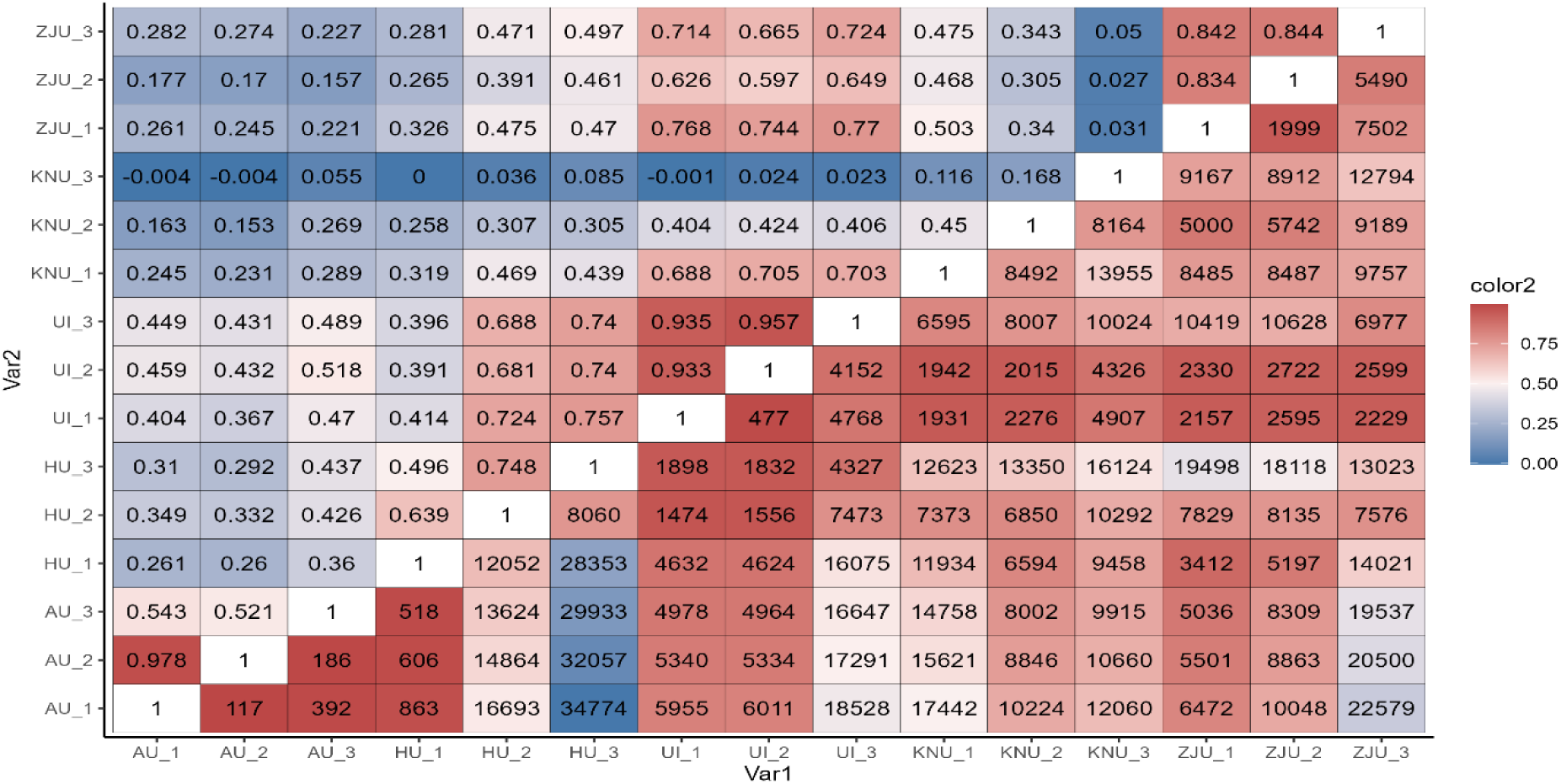
Association metrics between pairs of environments based on phenotypic information. The upper triangular matrix presents phenotypic correlations between location-by-year combinations. The lower triangular matrix contains the sum of the squares of the differences between the phenotypic observations from the same genotype observed in two different location-by-year combinations.

## Notes

### Competing Interest Statement

The authors have declared no competing interest.

